# Coupling transcriptomics and behaviour to unveil the olfactory system of *Spodoptera exigua* larvae

**DOI:** 10.1101/2020.05.22.110155

**Authors:** Angel Llopis-Giménez, Tamara Carrasco-Oltra, Emmanuelle Jacquin-Joly, Salvador Herrero, Cristina M. Crava

**Author notes:** Correspondence to: Salvador Herrero, Universitat de València. Department of Genetics, Dr Moliner 50, 46100 Burjassot, Spain, Cristina Crava, Universitat de València. Department of Genetics, Dr Moliner 50, 46100 Burjassot, Spain.

## Abstract

Chemoreception in insects is crucial for many aspects related to food seeking, enemy avoidance, and reproduction. Different families of receptors and binding proteins interact with chemical stimuli, including odorant receptors (ORs), ionotropic receptors (IRs), gustatory receptors (GRs), odorant binding proteins (OBPs) and chemosensory proteins (CSPs). In this work, we describe the chemosensory-related gene repertoire of the worldwide spread pest *Spodoptera exigua* (Lepidoptera: Noctuide) focusing on the transcripts expressed in larvae, which feed on many horticultural crops producing yield losses. A comprehensive *de novo* assembly that includes reads from chemosensory organs of larvae and adults, and other larval tissues, enabled us to annotate 200 candidate chemosensory-related genes encoding 63 ORs, 28 IRs, 38 GRs, 48 OBPs and 23 CSPs. Of them, 51 transcripts are new annotations. RNA-seq and reverse transcription PCR analyses show that 50 ORs are expressed in larval heads, and 15 OBPs are larva-specific. To identify candidate ecologically-relevant odours for *S. exigua* larvae, we set up behavioural experiments with different volatile organic compounds (VOCs). 1-hexanol triggers attraction at the three timepoints tested and linalool repels larvae at any timepoints. Other five VOCs elicit behavioural response at single timepoint. Lastly, we tested if pre-exposure to single VOCs influence the expression patterns of selected ORs and pheromone binding proteins (PBPs), showing a massive and general up-regulation of some ORs after 24h exposure. This work sets the basis for the study of chemosensation in *S. exigua* larvae, boosting further studies aimed to characterize chemosensory-related genes that underlie ecologically-relevant behaviours of larval stage.

## INTRODUCTION

*Spodoptera exigua* (Hübner, 1808) (Lepidoptera: Noctuidae), also known as the beet armyworm, is a worldwide spread Lepidoptera species. It is considered one of the most aggressive horticultural pests because of its high polyphagy, which provokes important economic losses (Zheng et al., 2011). The larval stage is the responsible for crop damage, since caterpillars feed on both the foliage and the fruits of different host plants (Greenberg et al., 2006).

Chemosensation is fundamental in shaping insects’ behaviours related to survival and reproduction, such as food searching, choice of oviposition substrate, mating seeking and detecting dangers like predators or parasitoids (Depetris-Chauvin et al., 2015; Robertson, 2015). In holometabolous insects such as Lepidoptera, the larval stage is devoted to food ingestion and growth whereas adults are dedicated to reproductive tasks. Larvae evaluate chemical cues from their ecological niche differently than adults, and the physiological and molecular equipment required for detecting odours is different between these two life stages (Dweck et al., 2018; Scherer et al., 2003).

Proteins involved in peripheral chemosensation include chemoreceptors and binding proteins. Volatile compounds are detected by odorant receptors (ORs) and antennal-expressed ionotropic receptors (aIRs) (Gomez-Diaz et al., 2018). ORs are seven-transmembrane domain proteins that work as heteromeric ligand-gated ion channels together with the odorant coreceptor (ORco) (Joseph and Carlson, 2017). Ionotropic receptors (IRs) are a divergent lineage of synaptic ionotropic glutamate receptors (iGluRs) (Rytz et al., 2013). Members of these two families are expressed in the dendrites of olfactory receptor neurons (ORNs), which stretch inside hair-like structures called olfactory sensilla. The olfactory sensillum surface has many pores where odorants can pass and activate the receptors in the ORNs, which transmit the signal to the higher brain centres. Olfactory sensilla are mainly located on the antenna and the maxillary palps. Non-volatile chemicals are sensed by gustatory receptor neurons (GRNs), which express the gustatory receptors (GRs) and a subset of IRs, the divergent IRs (dIRs) (Croset et al., 2010; Koh et al., 2015). GRNs are housed in gustatory sensilla, which are present in diverse body parts such as the mouth, the proboscis, the legs and even in the wings of the insects and are activated by direct contact with the chemical stimuli, which enter into the gustatory sensilla though an apical pore (Joseph and Carlson, 2017). Besides receptors, both olfactory and gustatory sensilla express two families of small soluble binding proteins which are secreted in the sensillar lymph: the odorant-binding proteins (OBPs) and the chemosensory proteins (CSPs) (Pelosi et al., 2005). OBPs are thought to carry the odorant molecules through the antenna lumen to the different receptors, but other functions have been proposed like odorant cleaning after its detection, protection of odorants from degradative enzymes and filtering of odorants (Sun et al., 2018; Zhou, 2010). Pheromone-binding proteins (PBP) are specialized OBPs that bind pheromone molecules, which are important for the insect mate recognition (Chang et al., 2015). Chemosensory proteins (CSPs) are soluble proteins also secreted in the sensillar lymph and, although their function is not clearly understood, they may play a role connecting the odorant molecules with the receptors (Pelosi et al., 2005).

Despite the larval stage of *S. exigua* is responsible for plant damage, there is a knowledge gap of the molecular machinery underlying larval olfaction and olfactory-driven behaviour. So far, many candidate chemosensory-related genes of *S. exigua* have been identified by RNA-seq analyses exclusively in adult tissues. Du *et al.* (2018) reported 157 candidate chemosensory genes identified in adult antennae, whereas Zhang *et al*. (2018) reported 159 ones identified in adult antennae, proboscis and labial palps. Unfortunately, both studies used a different annotation nomenclature, making difficult the comparison between sequences.

In this study, we aim to fill the gap on the knowledge of *S. exigua* larval olfaction through the analysis of a comprehensive RNA-seq dataset that includes several larval tissues. We expand the number of putative chemosensory-related genes described in *S. exigua* and propose a unifying gene nomenclature based on reconstructed phylogenetic trees and following names used in the annotated *Spodoptera frugiperda* genome (Gouin et al., 2017). As previous exposure to volatile organic compounds (VOCs) has been shown to influence OR gene expression in *S. exigua* adults (Wan et al., 2015), we also examined whether such a regulation occurs in *S. exigua* larvae. We show that some ORs are strongly up-regulated after being exposed to any of the odorants tested, irrespectively of the larval behaviour they trigger. Altogether, our results provide novel insights on the molecular basis of *S. exigua* larvae olfactory detection and behaviour.

## MATERIALS AND METHODS

### Insects

The *S. exigua* colony (SUI) used for all the experiments has been reared at University of Valencia on artificial diet (Bell and Joachim, 1976) at 25 ± 3 °C with 70 ± 5% relative humidity, using a photoperiod of 16:8 h (light:dark).

### RNA extraction, library preparation and sequencing

*S. exigua* fourth-instar larvae were dissected with a scalpel and heads, midgut and fat body were excised and homogenised in Trizol (Roche). Adult antennae (male and female), brains (male and female) and ovaries were excised from adults and homogenised in Trizol. Total RNA was extracted following Trizol manufacturer’s instructions. A second purification step was carried out using the RNeasy Mini Kit (Qiagen). Three replicates consisting of tissues excised from 16 larvae each were done for larval head, adult antenna (male and female each) and adult brain (male and female each). Only one replicate was prepared for ovaries, larval midgut and larval fat body. Library preparation and Ilumina Hiseq 2000 sequencing were both carried out by Novogen Technology Co. Ltd. (China). Raw reads are available at NCBI SRA database (Project number PRJNA634227).

### De novo assembly and annotation of chemosensory-related genes

Paired-end (PE) raw reads were trimmed and used for *de novo* assembly using Trinity v2.3.1 (Grabherr et al., 2013) with --min_kmer_cov 2 parameter. Trinity-assembled contigs were further clustered with Corset (Davidson and Oshlack, 2014) and the longest transcript from each Corset cluster was selected to obtain the final assembly. *De novo* assembled transcriptome is available at NCBI TSA database with accession number PRJNA634227. Annotation of transcripts encoding odorant receptors (ORs), ionotropic receptors (IRs), gustatory receptors (GRs), odorant-binding proteins (OBPs) and chemosensory proteins (CSPs). was performed with iterative blast searches using the amino acid sequences predicted from the *S. frugiperda* genome (Gouin et al., 2017) as query. Selected contigs were manually inspected, their coding sequences were predicted using BioEdit and *S. frugiperda* orthologs as a master, and 5’ and 3’ UTRs were removed. In few cases, when two contigs were overlapping and likely representing the same transcripts based on their alignments with *S. frugiperda* orthologous genes, they were merged together to create a consensus sequence. Redundant sequences were then identified by iterative generations of maximum-likelihood trees and removed from the dataset. Thus, the final dataset contains a non-redundant list of putatively unique transcripts, which likely correspond to unique genes, although we cannot exclude that some of them might be allelic variants of the same gene. Maximum-likelihood (ML) trees were built with protein sequences annotated from *S. frugiperda* (Gouin et al., 2017), *S. litura* (Zhu et al., 2018) and *B. mori* genomes (Forêt et al., 2007; Tanaka et al., 2009; van Schooten et al., 2016; Vogt et al., 2015; Wanner and Robertson, 2008), as well as putative proteins deduced from *S. littoralis* (Walker et al., 2019), *S. litura* (Gu et al., 2015), *Helicoverpa armigera*, *H. assulta* (Chang et al., 2017) and previous *S. exigua* transcriptomes (Du et al., 2018; Zhang et al., 2018). Trees were built using RAxML (Stamatakis, 2014) and amino acid MUSCLE alignments (Edgar, 2004) generated by MEGAX (Kumar et al., 2018). Based on the phylogenetic relationships, *S. exigua* chemosensory-related proteins were named according to the *S. frugiperda* nomenclature. Web based blastx searches were then run for *S. exigua* chemosensory-related transcripts whose corresponding proteins had no one-to-one ortholog in any *Spodoptera* species in order to verify their annotation as chemosensory-related genes.

### RNA-seq quantification of chemosensory-related gene expression

Expression levels of *S. exigua* chemosensory-related transcripts in larva head, female adult antenna and male adult antenna (3 replicates for each tissue) were estimated by mapping the trimmed reads to the chemosensory-related genes annotated in this study. Mapping was performed using Bowtie 2 (version 2.3.5.1) (Langmead and Salzberg, 2012) and RSEM (version 1.3.1) (Li and Dewey, 2011) with default parameters. Relative abundance of each candidate transcript is reported as TPM (Transcript per Million). For each transcript family, expression data were clustered by hierarchical clustering analysis using the heatmap.2 function from gplots v3.0.1.1 package of R software. Differential expression analysis between male and female antennae was carried out using EdgeR (Robinson et al., 2009). Transcripts were considered differentially expressed (DE) at false discovery rate (FDR) threshold < 0.05 and 2-fold change cut-off.

### Expression analysis of chemosensory-related transcripts by reverse transcription (RT)-PCR

Presence of transcripts encoding ORs, PBPs, two general OBPs (GOBPs) and three GR candidates for CO_2_ reception were analysed by RT-PCR in larva head and adult antenna in order to confirm their developmental expression specificity. The same RNA samples used for the RNA-seq sequencing (Llopis-Giménez et al., 2019) were employed. RNA pools were prepared for each of the two developmental stages, mixing the same amount of total RNA from each replicate (3 replicates from the larvae head, 6 replicates from the adult antenna). A total of 2 μg of RNA were treated with DNAseI (ThermoFischer Scientific) and converted into cDNA using PrimeScript cDNA synthesis kit (Takara), following the manufacturer’s protocols. Amplifications were run in a StepOnePlus Real-Time PCR system (Applied Biosystems) using 5x HOT FIREpol EvaGreen qPCR Mix Plus (ROX) from Solis BioDyne. The total reaction volume was 20 μl. Forward and reverse primers for all the transcripts were designed using the online software tool Primer3Plus (Untergasser et al., 2007). A list of primers used in this experiment is provided in Supplementary Table 1. RT-PCR products were run in a 2% agarose gel to visualize the amplification of a single band of the expected size.

### Chemicals

Chemicals used in this study were purchased from Sigma-Aldrich (95-99% purity) with the exception of methanol that was purchased from Labkem: propiophenone (CAS #: 93-55-0), cinnamaldehyde (CAS #: 104-55-2), cis-3-hexenyl propionate (CAS #: 33467-72-2), 3-octanone (CAS #: 106-68-3), trans-2-hexen-1-al (CAS #: 6728-26-3), benzaldehyde (CAS #: 100-52-7), cis-3-hexenyl acetate (CAS #: 3681-71-8), linalool (CAS #: 78-70-6), benzyl alcohol (CAS #: 100-51-6), hexyl propionate (CAS #: 2445-76-3), acetophenone (CAS #: 98-86-2), indole (CAS #: 120-72-9) and 1-hexanol (CAS #: 111-27-3).

### Behavioural assays

Behavioural assays were performed to study the effect (attraction or repellence) of VOCs to *S. exigua* larvae in a complex background. Ten fifth-instar *S. exigua* larvae were placed in one side of a 14 cm diameter Petri dish whereas a piece of artificial diet (1.5 × 0.8 × 1 cm) was placed at the opposite side. The Petri dish was placed inside a paperboard box (30 × 22 × 22 cm). A hole in the side of the box (6 cm of diameter) was made to include a 50 W halogen artificial light (at 15 cm of distance to the Petri dish). Fifty μl of the odorant diluted at 100 mg/ml in methanol were added to the artificial diet. In parallel, a control without odorant was run. Each odorant was tested a total of nine times. Replicates consisted of 3 biological replicates that used 3 different batches of larvae (*i.e*. larvae deriving from a different offspring). For each replicate, larval mobility was scored dividing the Petri dish in 10 areas of 1.3 cm each (Figure 1). To each area, we assigned a score from 0 to 9, which increased accordingly to the distance from the starting point. At time 2’, 5’ and 10’, the number of larvae in each area was recorded and the mobility index was determined as the sum of the scores obtained by each of the 10 larvae. The larval attraction index was calculated dividing the mobility index in presence of the odorant by that in the parallel control run. Values higher than 1 meant that the larvae were more attracted to the diet + odorant source than diet only, and values lower than 1 meant that larvae were less attracted to diet + odorant source than diet only (deterrent effect). Statistical analyses were conducted using a one-sample *t*-test comparing the attraction index at each time point with the theoretical value of 1. All statistical analyses were performed using GraphPad Prism software (v.7.0).

**Figure 1.**
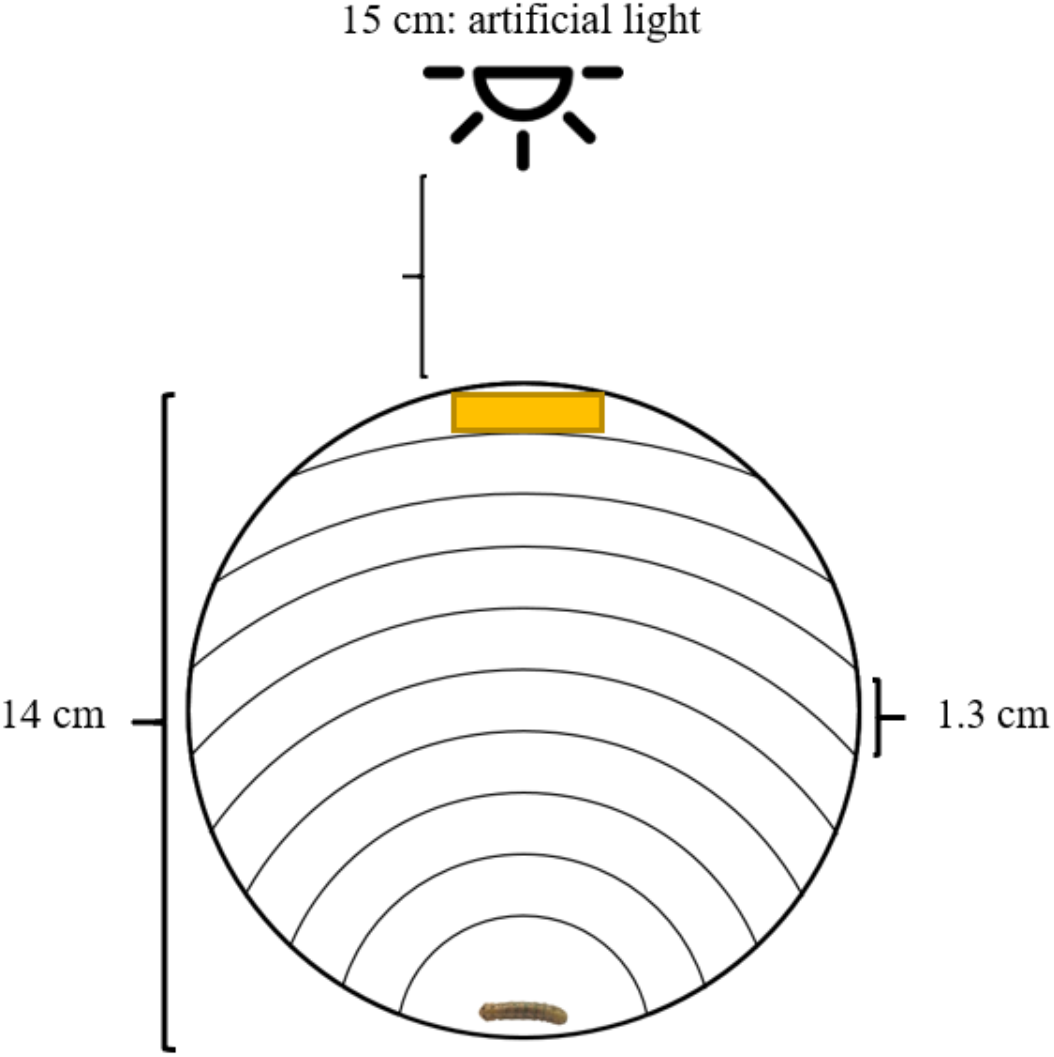
Scheme of the behaviour assay design. Ten 5^th^-instar larvae were put in one of the sides of the Petri dish whereas a piece of artificial diet was placed on the opposite side. A halogen artificial light was positioned at 15 cm from the Petri dish. Five mg of the odorant (or the equivalent volume of solvent) were placed on the artificial diet.

### Odorant exposure and tissue collection

To test the influence of VOC exposure on the transcriptional profile of ORs and OBPs, twenty *S. exigua* fourth-instar larvae were placed in a 9 cm Petri dish including inside a perforated 1.5 ml tube containing a Whatmann paper soaked with 50 μl of the odorants (100 mg/ml). Control consisted of exposure to methanol solvent only. Petri dishes were kept at 25°C. Ten larvae heads were dissected at 1h and 24h of exposure and stored in 300 μl of Trizol Reagent (Invitrogen) at −80°C for RNA extraction. Three independent replicates for each treatment were done.

### Starvation and tissue collection

The transcript levels of ORs and OBPs were also measured under starving conditions. Sixteen *S. exigua* fourth-instar larvae were let at 25 °C without any food for 24 h, whereas control larvae were allowed to feed on artificial diet. Larvae heads were dissected after 24 h and stored in 300 μl of Trizol Reagent (Invitrogen) at −80°C for RNA extraction. Three independent replicates for each treatment were done.

### RNA extraction, cDNA synthesis and quantitative real-time PCR (RT-qPCR)

Total RNA was purified using Trizol reagent following the manufacturer’s instructions. 500 ng of each RNA was treated with DNAseI (ThermoFischer Scientific) following manufacturer’s protocol. Then, samples were converted into cDNA using SuperScript II Reverse Transcriptase (ThermoFischer Scientific) following manufacturer’s recommendations and using random hexamers and oligo (dT) primers. In order to help to the nucleic acid precipitation, 10 μl of Glycogene (Roche), were used per sample. RT-qPCR was performed in a StepOnePlus Real-time PCR system (Applied Biosystems) using 5x HOT FIREpol Eva Green qPCR Mix Plus (ROX) (Solis Biodyne) in a total reaction volume of 20 μl. Forward and reverse primers for every gene were designed using the online software tool Primer3Plus (Untergasser et al., 2007). An endogenous control *ATP synthase subunit C* housekeeping gene was used in each qPCR to normalize the RNA concentration. A list of the used primers is provided in the Supplementary Table 1. The differences in expression between treatments (control and infected) were calculated using the ∆∆Ct method (Livak and Schmittgen, 2001). A one-sample *t-*test was used to search for statistical differences comparing each Log Fold-Change (LogFC) value to the theoretical value of 1. Graphs and the statistical analysis were performed using GraphPad Prism software (v7.0). Heatmaps were performed using the R packages gplots and RColorBrewer.

## RESULTS

### Annotation of chemosensory-related genes

We provide in this study an updated repertoire of 200 candidate chemosensory-related genes belonging to 5 different families: 63 ORs, 28 IRs, 38 GRs, 48 OBPs and 23 CSPs, (Table 1). These results greatly expanded previous annotations of chemosensory-related genes in *S. exigua* since 51 genes appear to be newly annotated: 5 ORs, 5 IRs, 22 GRs, 16 OBPs and 3 CSPs (Figure 2, Supplementary Figure 1-4). Moreover, our annotation efforts provide a new phylogeny-based nomenclature that follows the one in *S. frugiperda* (Gouin et al., 2017) and in *S. littoralis* (Walker et al., 2019) with the aim to aid future comparative studies among related species.

**Table 1.** Summary of the candidate chemosensory-related genes of *Spodoptera exigua* annotated in this study.

| Gene family | *S. exigua* |  |  |  | *S. frugiperda*<sup>c</sup> |
| --- | --- | --- | --- | --- | --- |
|  | This study |  | Du's dataset<sup>a</sup> | Zhang's dataset<sup>b</sup> |  |
|  | All | New |  |  |  |
| OR | 63 | 5 | 50 | 64 | 69 |
| IR | 28 | 5 | 20 | 22 | 43 |
| GR | 38 | 22 | 7 | 30 | 233 |
| OBP | 48 | 16 | 45* | 24 | 51 |
| CSP | 23 | 3 | 32* | 19 | 22 |

<sup>a</sup>Du et al., (2018)
<sup>b</sup>Zhang et al. (2018)
<sup>c</sup>Gouin et al. (2017)
\* These numbers include several transcripts likely mis-annotated as *S. exigua* OBPs and CSPs (Supplementary table 3)

**Figure 2.**
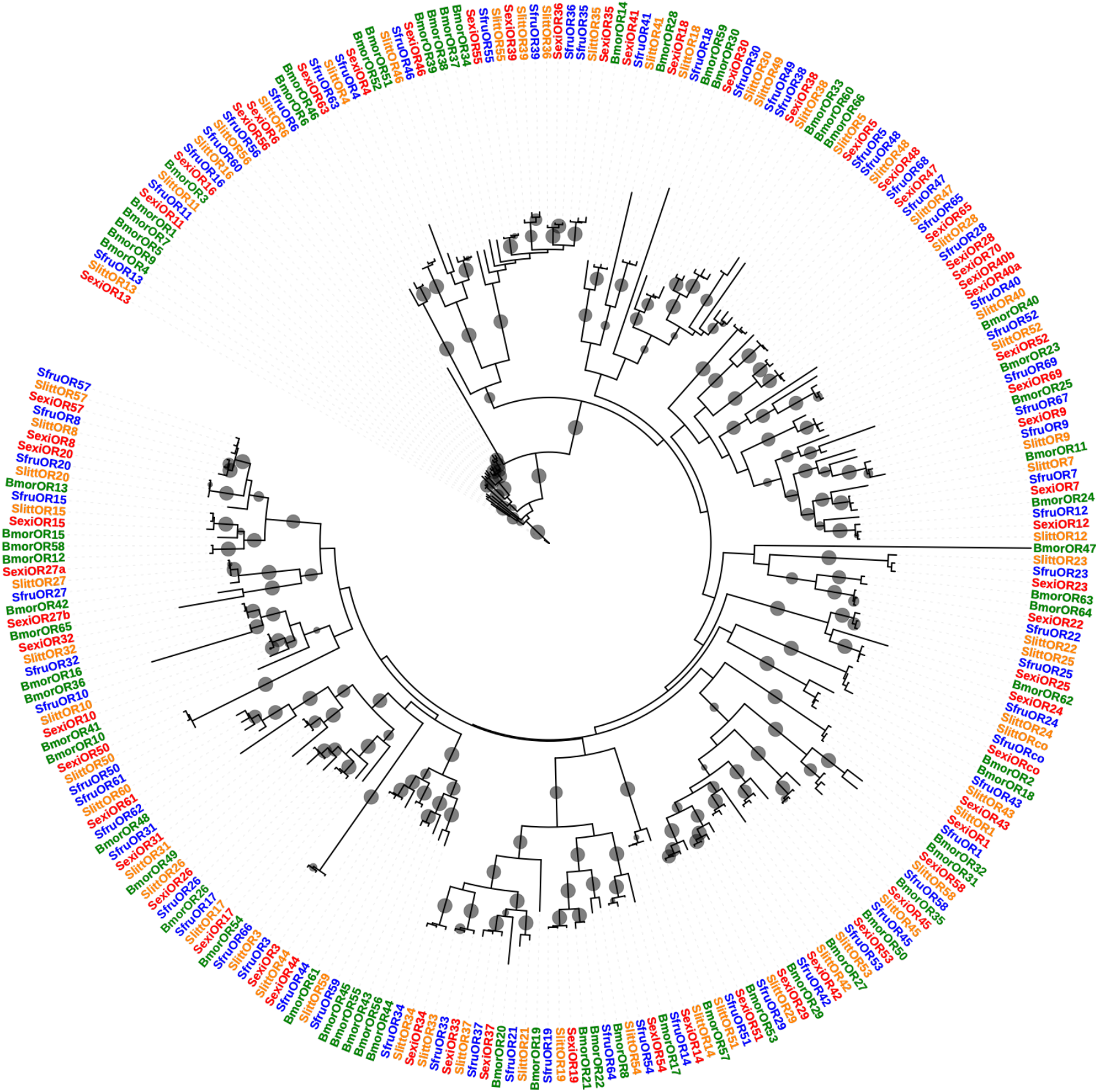
Phylogenetic tree of *Spodoptera exigua* odorant receptors (SexiORs). Maximum-likelihood (ML) tree built with protein sequences annotated from *S. frugiperda* (Gouin et al., 2017) and *B. mori* genomes (Tanaka et al., 2009) as well as putative proteins annotated from *S. littoralis* transcriptome (Walker et al., 2019). SexiORs are shown in red, *S. frugiperda* ORs in blue, *S. littoralis* ORs in yellow, and *B. mori* ORs in green. Grey dots show a bootstrap value higher than 80.

A total of 63 ORs have been annotated (Table 1). Of them, 44 (70 %) have complete ORFs. Five ORs have not been previous reported by any previous study (SexiOR46, SexiOR54, SexiOR56, SexiOR69 and SexiOr40b) (Supplementary Table 2). All *S. exigua* ORs (SexiORs) have a one-to-one ortholog relationship with *S. frugiperda* ORs except SexiOR40b and SexiOR40c that might represent OR40 lineage-specific duplications in *S. exigua*, although we cannot exclude they are transcriptional isoforms of the same gene (Figure 2). Our annotation effort missed three incomplete candidate SexiORs described by Zhang *et al.* (2018) (namely OR8, OR56, and OR59 according to Zhang’s nomenclature).

We identified 28 IRs in the *S. exigua* transcriptome (SexiIRs) (Table 1), and all of them have a one-to-one ortholog in *S. frugiperda* (Supplementary Figure 1). Of them 14 (50%) have a complete ORF. Compared to previous works in *S. exigua*, five IRs have been newly annotated (namely SexiIR7d.2, SexiIR100a, SexiIR100b, SexiIR100c and SexiIR100i) (Supplementary Table 2). All of the newly annotated genes belong to the divergent IRs subfamily (Guo et al., 2017).

Thirty-eight candidate *S. exigua* GRs (SexiGRs) were found in our transcriptome (Table 1) but only ten sequences were complete (26%) (Supplementary Table 2), likely due to the low levels of expression commonly described for GRs (Dunipace et al., 2001). Orthologs to GR1, GR2 and GR3 (Guo et al., 2017), the antenna-expressed and well-conserved CO_2_ receptors, were found completed (Supplementary Figure 3). Twenty-two GRs were newly described compared to previous annotations in *S. exigua* (Supplementary Table 2). Identification of orthologous genes in *S. frugiperda* has been difficult due to the incomplete ORF retrieved for the majority of SexiGRs, which led to low branch support in many cases (Supplementary Figure 2). Compared to the annotation made by Zhang *et al.* (2018), 11 of the GRs annotated in their study were not present in our transcriptome. Special mention should be made to GR24 (according to Zhang’s nomenclature), also mis-annotated by Du et al. (2018) as the CO_2_ receptor GR1. Both annotations report a sequence of a likely partial GR (136 amino acids for Zhang’s GR24 and 120 for Du’s GR1). We retrieved the same sequence in our transcriptome, but blast searches and phylogenetical analysis clearly showed that it corresponds to an unrelated and uncharacterized protein of around 120 amino acids presents in the genomes of Lepidoptera species (Supplementary Figure 3).

Forty-eight *S. exigua* OBP candidate transcripts were identified (SexiOBPs) (Table 1). Most of them are complete (69%), and only 15 had a partial ORF (Supplementary Table 2). Sixteen transcripts were newly annotated. All SexiOBPs have a clear one-to-one orthologue in *S. frugiperda* except SexiOBP46 and SexiOBP47 (Supplementary Figure 4). Two of the transcripts described by Zhang *et al.* (2018) (OBP25 and OBPN-3) were not identified in our transcriptome. Compared to Du *et al.* (2018), our annotation missed 19 putative OBPs. However, a closer look to these missing OBPs revealed that they did not cluster with any *Spodoptera* OBPs. Twelve of them grouped with *Helicoverpa* OBPs in our phylogenetic tree whereas the other seven were far distant (Supplementary Figure 4). Further analysis with blastx revealed that the best hit of missing OBPs was against sequences from distantly-related species (such as butterflies or Coleoptera species) (Supplementary Table 4). Consequently, we suspect that these nineteen sequences (which were neither annotated by Zhang et al., 2018) might arose from contamination during library preparation and sequencing.

Twenty-three candidate *S. exigua* CSP transcripts have been annotated (SexiCSPs) (Table 1). Of them, 21 have a complete ORF (91%). Three of the annotated sequences have not been described in any prior study: SexiCSP11, SexiCSP23 and SexiCSP24 (Table 1 and Supplementary Table 2). All SexiCSPs have a clear one-to-one ortholog in *S. frugiperda* except SexiCSP23 and SexiCSP24 (Supplementary Figure 5). Our annotation retrieved all but one CSP (CSP-N3) described Zhang *et al.* (2018). Compared to annotation reported by Du et al (2018) we missed many tentative CSPs (fifteen) that were also absent in Zhang’s annotation. However, likewise missing OBPs, missing CSPs likely arose from sample contamination. Eight of them clustered with *Helicoverpa* sequences instead of *Spodoptera* ones and the remaining sequences had best blast hit against sequences from distantly-related Lepidoptera species or from other insect orders (Coleoptera, Diptera and Hymenoptera).

### Chemosensory-related transcripts in S. exigua adults and larvae

Transcript levels were evaluated mapping RNA-seq reads obtained from larval head, female and male antennae to our manually curated chemosensory-related gene dataset. Since data from larvae and adults were not directly comparable because they proceeded from different tissues (isolated antennae in adults versus whole heads from larvae), we could not relate expression levels between adults and larvae. Instead, we used these data to 1) establish the larval chemosensory-related gene set by detecting the ones that had transcription signals in larval head samples, 2) identify which genes were differentially expressed between male and female antennae.

Mapping of reads obtained from larval head to the manually curated chemosensory-related transcript dataset only gave us a hint about expressed genes without providing any information of transcripts not expressed at all, since RNA-seq data from composite structures (such as head) often lead to false negatives (Johnson et al., 2013). In total we found 30 ORs, 22 IRs, 13 GRs, 35 OBPs and 20 CSPs that had mapping reads in at least one of the larval head replicates (Figure 3, Supplementary Figures 7 – 10). By qualitative comparison with the adult chemosensory-related gene set, we noticed that the larval gene set was smaller than that of adults (120 versus 187). We found a larval-specific expression only for OBPs: 14 OBPs had mapping reads from larval heads but not from adult antennae (Supplementary Figure 9) and 8 of them had been newly annotated. RNA-seq data have been confirmed using RT-PCR for ORs, putative CO_2_ receptors (SexiGR1, SexiGr2 and SexiGR3) and some OBPs whose function was previously characterized (Liu et al., 2015a). Our results showed that 50 out of 63 ORs were actively transcribed in larval heads, whereas all ORs were detected in adult antennae tissue, thus revealing that 14 ORs were adult-specific (*i.e.* SexiOR1, SexiOR7, SexiOR10, SexiOR17, SexiOR20, SexiOR24, SexiOR26, SexiOR34, SexiOR37, SexiOR38, SexiOR40a, SexiOR43 and SexiOR44). GOBPs and PBPs appeared to be expressed in both larval heads and adult antennal tissues. Of three candidate CO_2_ GRs, our results only SexiGr1 and SexiGR2 are expressed in larval heads and adult antennae whereas SexiGR3 was not expressed in any sample.

**Figure 3.**
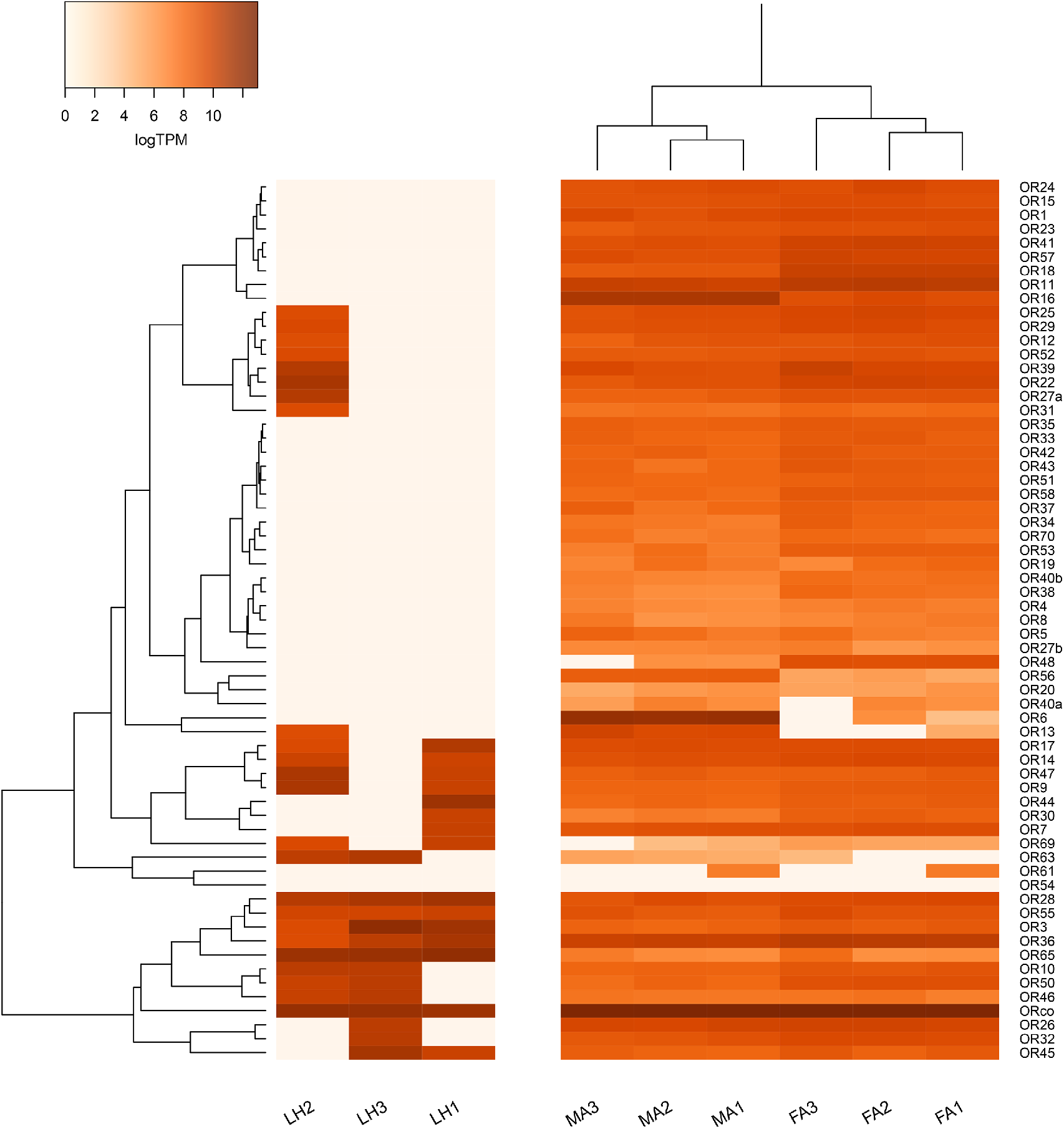
Heat-plot of odorant receptors (SexiORs) expression in whole head of *Spodoptera exigua* larvae and adult antennae. Colour plots represent log2 of transcripts per million (TPM) values estimated by RSEM. LH: Larvae Head. MA: Male Antenna. FA: Female Antenna. Asterisks indicate statistically significant differences between male and female antenna samples identified by EdgeR analysis (FDR<0.05).

Differentially expressed (DE) transcripts between male and female adults were 17. Of these, ten had significantly higher expression in male antenna and seven in female antenna. Transcripts with higher expression in males were four ORs, four OBPs, one GR and one CSP. Among the DE transcripts that showed the highest variation by differences in expression (more than 4-fold change) there were ORs and OBPs involved in pheromone binding (SexiOR6, SexiOR13, SexiOR16 and SexiPBP1). Transcripts upregulated in females had less variation by differences in expression than transcripts upregulated in males. In female antennae, fold-changes varied from 2.3 to 3, except for SexiOR48 that was 20-fold more expressed in female than in male antenna.

### Behavioural experiments

The behavioural response of *S. exigua* 5^th^ instar larvae to thirteen volatile compounds was investigated (Figure 4). 1-hexanol and benzaldehyde were the only tested volatiles that enhanced artificial diet attractancy whereas five other compounds had a deterrent effect. 1-hexanol evoked enhanced attraction at the three time-points whereas benzaldehyde was active only at one time point. Indole exhibited a deterrent effect at the three tested time-points, 3-octanone at both 5’ and 10’ time-points, benzyl alcohol at the 10’ time-point only, linalool and cis-3-hexenyl propionate at the 5’ time-point only. The remaining odorants (propiophenone, cinnamaldehyde, trans-2-hexen-1-al, cis-3-hexenyl acetate, hexyl propionate and acetophenone) did not show any significant effect on diet attraction at any time.

**Figure 4.**
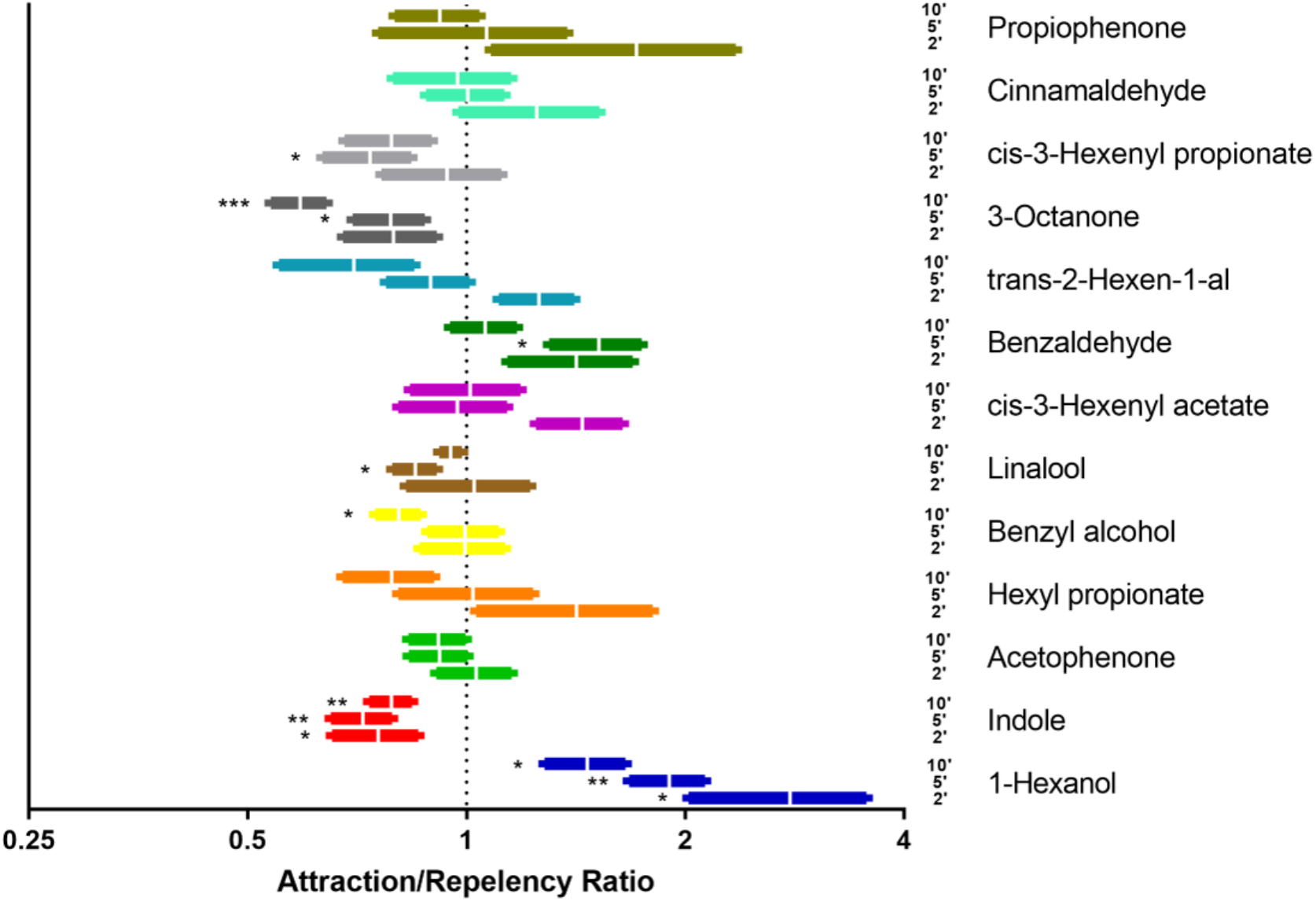
Larval attraction index of *Spodoptera exigua* larvae to different odorant stimuli at different times. Fifty μl diluted at 100 mg/ml of each odorant were used for the bioassay. Bars represent the mean value and the standard deviation. Values above 1 are indicative of attraction and values below 1 are indicative of repellence. Asterisks indicate statistically significant differences (one-sample *t*-test) (P < 0.05 *, P < 0.01 **, P < 0.001 ***).

### Regulation of larval chemosensory-related gene expression after odorant exposure

Expression levels of selected ORs and PBPs expressed in larvae were analysed by RT-qPCR after VOC exposure. We observed that short-time exposure (1h) triggered low levels of variation of few ORs to some specific VOCs. 1-hexanol exposure increased the expression of SexiOR23 and SexiORco (2.5-fold change and 2.3-fold change, respectively). Indole exposure up-regulated 20.9-fold the expression of SexiOR25 and acetophenone increased 5.8-fold the expression of SexiOR11. Benzaldehyde down-regulated 1.4-fold the expression of SexiOR65 (Figure 5). Cis-3-hexenyl acetate exposure did not induce any significant variation.

**Figure 5.**
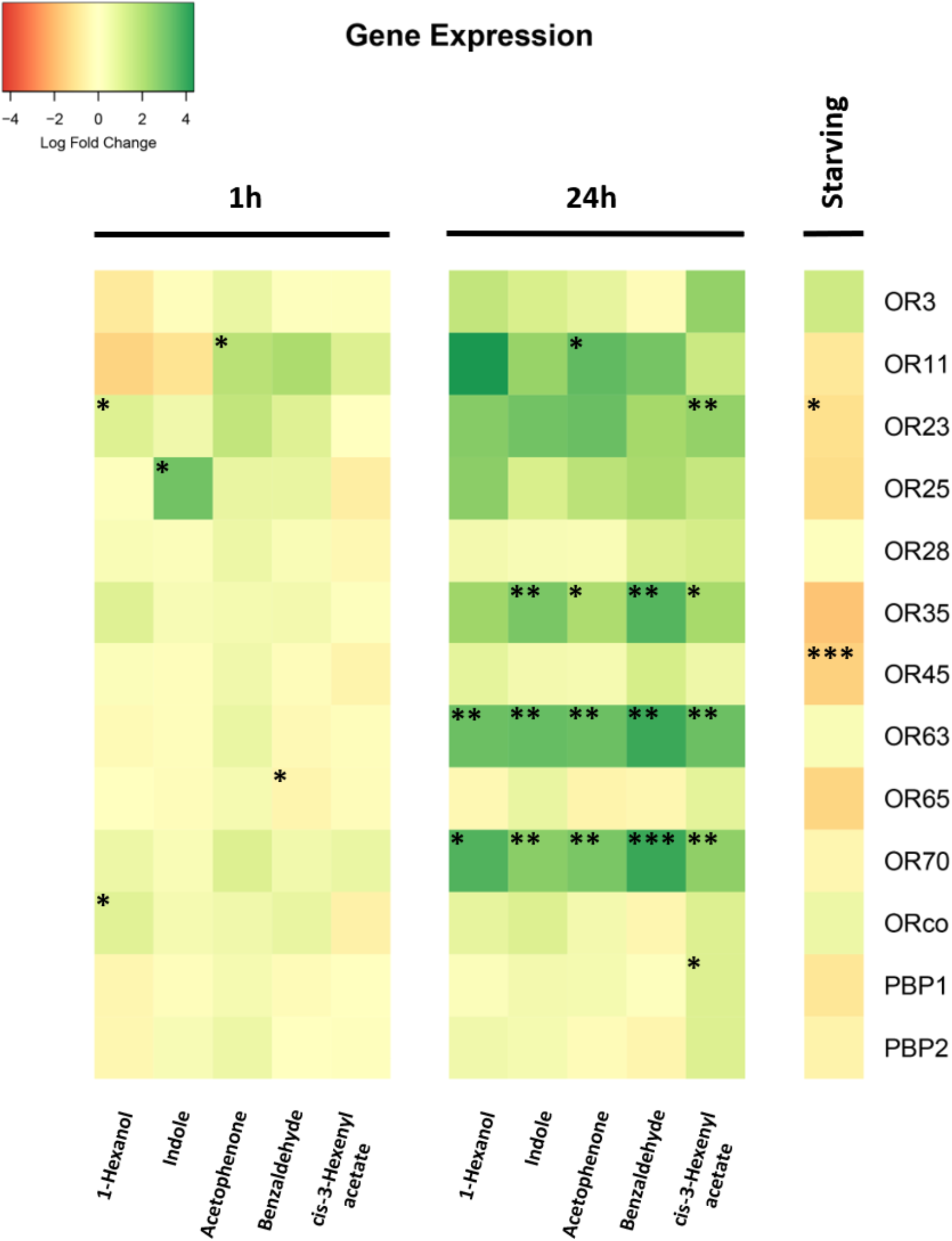
Heat-plot of relative expression levels of odorant receptors (SexiORs) and pheromone-binding proteins (SexiPBPs) in *Spodoptera exigua* larvae after exposure to different odorants or in starving conditions. Expression values were estimated by RT-qPCR using the ∆∆Ct method. Colour plots represent Log fold change values. Dark red colours indicate a decrease in the expression and dark green ones indicate an increase in the expression. Asterisks indicate statistically significant differences (one-sample *t*-test) (P < 0.05 *, P < 0.01 **, P < 0.001 ***).

Long-term exposure (24h) led to wider transcriptional changes than short-time exposure. All the odorants tested induced expression changes of 2 to 5 genes, depending on the odorant. Interestingly, exposure to any of the tested odorants only triggered strong up-regulation of mRNA levels and never down-regulation. Expression of SexiOR63 and SexiOR40c, was strongly up-regulated after exposure with any of the odorants tested (from 12- to 49-fold changes). The mRNA levels of SexiOR35 increased after exposure to any odorants although only the changes triggered by four out of the five odorants were statistically significant (fold changes varied from 7- to 33-fold). Likewise, expression of SexiOR11, SexiOR23 and SexiOR25 showed a general trend of up-regulation after exposure to any odorants, although only SexiOR11 changes after acetophenone exposure (27.7-fold) and SexiOR23 changes after cis-3-hexenyl acetate (11.7-fold) were statistically significant. This last compound also triggered SexiPBP1 up-regulation (2.6-fold change) (Figure 5).

### Regulation of larval chemosensory-related genes under starvation

The same set of genes whose expression was tested after odorant exposure was also used to analyse mRNA levels after starvation. The absence of food ingestion for 24h led to down-regulation of only SexiOR23 (2.8-fold) and SexiOR45 (4.1-fold) (Figure 5).

## DISCUSSION

In Lepidoptera, the study of olfaction has been mainly limited to adult stage, and focused on the understanding of sex-linked behaviours such as sex pheromone detection and egg-laying substrate selection (Allison and Carde, 2016; García-Robledo and Horvitz, 2012; Haverkamp et al., 2018). In contrast, the molecular and physiological mechanisms that underlie olfactory behaviours in larvae are poorly understood and only few reports have described the chemosensory gene set expressed at the larval stage (Chang et al., 2017; Di et al., 2017; McCormick et al., 2017; Poivet et al., 2013; Tanaka et al., 2009; Walker et al., 2016) compared to the plethora of adult transcriptomes available (Montagné et al., 2015). Here, we identified chemosensory-related genes expressed in *S. exigua* larval heads using RNA-seq and RT-PCR data, expanding previous datasets built from adult data (Du et al., 2018; Zhang et al., 2018). We also developed a method to analyse *S. exigua* larval response to volatiles and we demonstrated that exposure to selected odorants drives changes in OR transcription.

Previous efforts for annotation of chemosensory-related genes in *S. exigua* reported 157 and 159 candidate transcripts identified in adult tissues (Du et al., 2018; Zhang et al., 2018). Our results greatly expanded these previous annotations since we report 200 chemosensory-related genes annotated from RNA-seq from multiple adult and larval tissues. Moreover, our annotation identified mis-annotated transcripts previously reported as *S. exigua* candidate GRs, OBPs and CSPs, and provided a more reliable nomenclature of *S. exigua* chemosensory-related candidate genes based on orthologs identified in the *S. frugiperda* genome (Gouin et al., 2017). In total, we identified 63 ORs, 28 IRs, 38 GRs, 48 OBPs and 23 CSPs. *S. frugiperda* and *S. litura* are two species closely related to *S. exigua* whose genomes were fully sequenced (Cheng et al., 2017; Gouin et al., 2017). The number of ORs in both genomes ranges from 69 to 73, suggesting that the ORs described in *S. exigua* almost cover the full OR repertoire in this species. OBP and CSP repertoires are likewise almost complete since these two gene families have 51 and 22 members in *S. frugiperda*, and 36 and 23 members in *S. litura*, respectively. On the contrary, the number of GRs is far lower than what has been described in other *Spodoptera* spp. from genome analyses (231 GRs in *S. frugiperda* and 237 in *S. litura*), likely due to the sampled tissues and the low expression level of GRs (Dunipace et al., 2001). Many IRs members are still missing for *S. exigua* compared to the 43 IRs present in *S. frugiperda* genome. In Lepidoptera, this gene family is divided in three subclasses: antennal IRs (aIRs), divergent IRs (dIRs) and Lepidoptera-specific IRs (lsIRs) (Liu et al., 2018). Of these, aIRs are highly conserved in sequence and in Lepidoptera they clustered in 16 orthologous groups that are largely characterized by a one-to-one orthologous relationship. Here, we describe the homologous sequence of the 17 aIRs present in *S. frugiperda* genome (Gouin et al., 2017; Liu et al., 2018). Hence, the missing *S. exigua* IRs probably belong to the dIRs and lsIRs subclasses, which are characterized by lineage-specific expansions (Liu et al., 2018).

Expression data for ORs and other chemosensory-related transcripts in *S. exigua* have been limited to the main adult olfactory tissues, antenna and maxillary palps (Du et al., 2018; Liu et al., 2015b; Wan et al., 2015; Zhang et al., 2018). Here we provide the first comprehensive dataset of chemosensory-related transcripts expressed in the larval stage. We combined RNA-seq and RT-PCR as RNA-seq data from composite tissues (such as head) often delivers false negative results (Johnson et al., 2013), especially for genes expressed in few cells and at a low level, which is the case of chemosensory receptors. Altogether, we found 50 out of 63 ORs expressed in larvae, including the four sex pheromone receptors (Liu et al., 2013). Notably we did not observe any larval specific OR. Similarly, no larval-specific ORs have been found in transcriptomic data from *S. littoralis* (although the number of larval expressed ORs was lower than what we observed in *S. exigua*: 22 out of 47), *Dendrolimus punctatus* and *Lymantria dispar* larvae (Poivet et al 2013, Zhang et al 2017, McCormick et al 2017). On the contrary, six larval-specific ORs have been identified in *Bombyx mori* (Tanaka et al., 2009), one in *H. armigera* (Di et al., 2017) and one in *C. pomonella* (Walker et al., 2016).

Contrary to ORs, we observed 14 larval-specific OBPs in *S. exigua*, and we can speculate they are involved in binding specific cues detected by larvae. One larval-specific OBPs has been found in *S. littoralis* (Poivet et al., 2013), and 10 were found in *L. dispar* (McCormick et al., 2017), suggesting that the occurrence of larval specific OBP might be common in Lepidoptera. We observed the expression of the four pheromone-binding proteins (PBPs) in *S. exigua* larval heads. PBPs and pheromone receptors are part of the molecular machinery used by adult moths to detect the sex pheromone and their expression in *S. exigua* larvae is intriguing. This, however, corroborates previous studies that reported PBP and pheromone receptor expression in larvae from diverse Lepidoptera species (Jin et al., 2015; Poivet et al., 2012; Zhu et al., 2016; Zielonka et al., 2016). As proposed by Poivet et al (2012), larvae may use the pheromonal signal to find food.

We have annotated the three putative Lepidoptera CO_2_ receptors in *S. exigua*, but only two of them were found to be expressed in larval head and adult antenna. Similarly, only two are expressed in adult antenna and proboscis in *S. littoralis* (Walker et al., 2019). It is probable that these two transcripts are sufficient for CO_2_ sensing in *S. exigua*, as it has been shown that only two out of the three CO_2_ receptors indispensable and sufficient for CO_2_ sensing (Ning et al., 2016, Xu et al., 2020).

Differential expression analysis between male and female adult antenna identified seventeen chemosensory-related transcripts with a sex-biased expression. As expected, we found among transcripts upregulated in males three sex pheromone receptors (SexiOR6, SexiOR13 and SexiOR16) and one pheromone-binding protein (SexiPBP1), which are involved in sexual communication (Liu et al., 2015a, 2013). Our results support those of Liu et al., (2013, 2015), which showed male-biased expression of these pheromone receptors and SexiPBP1, using RT-qPCR (Liu et al., 2015a, 2013). The fourth *S. exigua* candidate pheromone receptor, SexiOR11, did not exhibit differential expression between sexes in our study, nor in Liu et al (2013), but in contrast to Du et al (2018) study that found a male-biased expression for this OR. It has to be noticed that this OR is not yet confirmed as a pheromone receptor, since it failed to respond to any pheromone tested (Liu et al 2013). In addition to pheromone related transcripts, we noticed another OR with a male-biased expression, SexiOR56. Together with the fact that this OR clustered in the sex pheromone receptor clade of Lepidoptera, we suspect SexiOR56 may be a fifth pheromone receptor present in *Spodoptera* spp. In female antenna, all the seven female-enriched transcripts were ORs. We can speculate they may detect volatiles involved in female important behaviours such as egg-laying substrate searches and choice. Of them, only the two that showed the greatest variation (SexiOR18 and SexiOR48) matched with the results obtained by Du et al. (2018).

Larvae detect odours in the environment to succeed in many ecological tasks such as selecting host plants or moving to more palatable food sources in the same plant, escaping from parasitoids, detecting harmful microbes or correct places for pupating (Becher and Guerin, 2009; Carroll et al., 2008, 2006; Carroll and Berenbaum, 2002; Ebrahim et al., 2015; Mooney et al., 2009; Piesik et al., 2008; Poivet et al., 2012; Singh and Mullick, 2002; Stensmyr et al., 2012; Tanaka et al., 2009; Zhu et al., 2016). Identification of behaviourally active odorants is the first step needed to link volatile molecules to larval ecology. Yet, only data for pheromone-triggered behaviour are available for *S. exigua* larvae (Jin et al., 2015). Our study investigated the behavioural response of fifth instar *S. exigua* larvae to thirteen VOCs that are known to be commonly emitted by plants. Of these, 1-hexanol, a ubiquitous plant volatile, and benzaldehyde, the primary component of bitter almond oil, enhanced larvae attraction to artificial diet. 1-hexanol has been shown to be attractive for other Lepidoptera species. In the closely related species *S. littoralis*, 1 -hexanol was attractive to larvae when the odorant was presented alone (de Fouchier et al., 2018) and when it was presented together with artificial diet (Rharrabe et al., 2014). 1-hexanol was also attractive for the larvae of *Lobesia botrana* (Becher and Guerin, 2009). On the contrary, in the domesticated species *B. mori*, 1-hexanol did not show any attraction in behaviour assays (Tanaka et al., 2009). Benzaldehyde was less active than 1-hexanol in our essays. In *S. littoralis* larvae, this compound was attractive at high doses (10 and 100 μg) but repellent at 0.1 μg (de Fouchier et al., 2018). In *H. armigera* larvae, it does not have any effect (Di et al., 2017). We observed a significant attraction at 5’ that was lost at 10’. A general tendency we observed was that the odorant effect decreased with time, probably due to saturation of the odorant in the Petri dish after the first minutes.

Five odorants were identified as deterrents to fifth instar *S. exigua* larvae. The most consistent responses were obtained for indole and 3-octanone. This last volatile, in addition to linalool that also acted as a deterrent, have been identified as major components of frass volatiles produced by larvae of a Noctuidae species (*Pseudoplusia includens*) and are used by parasitoids to locate host presence (Ramachandran and Norris, 1991). Thus, *S. exigua* larvae might avoid spots with high concentrations of these volatiles that may betray the presence of competitor larvae and their presence to natural enemies. Indole was acting as a deterrent at all time-point tested. Indole has been shown to enhance susceptibility of *S. exigua* larvae to two common Lepidoptera pathogens: baculovirus and *Bacillus thuringiensis* (Gasmi et al., 2019). Larvae might avoid indole-rich feeding spots that might alter their susceptibility to pathogens. However, in *S. littoralis*, indole was slightly attractive (de Fouchier et al., 2018). Another odorant that has been shown to promote larval susceptibility to baculovirus and *B. thuringiensis* was linalool, whereas 3-cis-hexenyl acetate did not have any effect (Gasmi et al., 2019). Consistently, we observed that linalool was acting as a deterrent to *S. exigua* whereas cis-3-hexenyl acetate did not trigger any statistically significant effect. Among the other behaviourally active compounds on *S. exigua* larval, benzyl alcohol showed a deterrent effect at the last time-point whereas it was attractant to *S. littoralis* larvae in a dose-response manner (de Fouchier et al., 2018).

In *S. exigua* adults, previous exposition to sex pheromone or plant volatiles triggered a broad up-regulation of several ORs and OBPs, including some known pheromone-receptors (Wan et al., 2015). This has been explained as a likely mechanism that mediates odour sensitization (Dion et al., 2019), a phenomenon that has been observed in *S. littoralis* (Anderson et al., 2013). Here we tested if mRNA levels of a set of ORs and PBPs in larvae was influenced by previous exposure to some of the behaviourally-active odorants identified in this study. We observed few and specific changes in gene expression after 1h of exposure, among which that of SexiOR25 showed a 21-fold change after indole exposure. Twenty-four h after exposure, we observed up-regulation of several OR expression, whatever the behavioural effects of the odorants. This observation is similar to what has been noticed in *S. exigua* adults (Wan et al., 2015), but in this latter study all transcripts tested appeared as up-regulated upon pheromone or plant volatile exposure, including SexiORco and SexiPBP1. Here, we did not observe any change in the expression of these transcripts in larvae (except for a limited but statistically significant fold-change in SexiPBP1 expression after cis-3-hexenyl acetate exposure). Similarly, we did not observe any change in SexiOR3 expression, a receptor narrowly tuned to E-ß-farnesene (Liu et al., 2014), whereas Wan et al., (2015) found its expression to be modulated upon adult host plant exposure. A study conducted in mammals and *Drosophila melanogaster* reported that exposure to a high dose of a given odorant would trigger down-regulation, and not up-regulation, of the expression of the responding OR, suggesting exposure experiment could be used to identify OR-ligand pairs (Von Der Weid et al., 2015). However, a thorough screening in *Drosophila* revealed in fact that OR expression increased, decreased or did not change upon exposure to the OR ligand, depending on the OR-ligand pair (Koerte et al., 2018). Here, we observed only up-regulation, whatever the odorant. It has to be noticed that we exposed insects to odorants for 24h, which is much more than the 5h exposure used in these former studies. Because the ligands for the ORs investigated here are not known (expect for SexiOR3 that was not regulated upon exposure to its ligand), it is thus difficult to interpret if transcription regulation of ORs is trigger by odorant exposure or if, alternatively, the observed up-regulation is a general stress response due to exposure to very high odorant doses during a long time. To test this latter hypothesis, we challenged the larvae to another stress, which consisted of 24h starvation. Upon starvation, we recorded a different effect: only two ORs were down-regulated. A first step to answer whether the observed OR up-regulation is due to stress or to specific induced transcription, functional studies are needed to identify if up-regulated ORs are indeed involved in the detection of the corresponding odorants.

In conclusion, our work provides i) a reliable annotation of the chemosensory-related transcripts in the noctuid pest *S. exigua*, focusing on larval expressed genes; ii) a new method to identify behaviourally-active VOCs against *S. exigua* larvae; iii) the evidence that long-term odorant exposure triggers broad changes in ORs expression. The data shown here represents the first step for further studies aimed to characterize receptors and their cognate ligands that are important for the ecology of *S. exigua* larvae. Identification of larval attractants or repellents has the additional value to be of further interest in the development of olfactory-based control techniques, which might help in protect crops from larvae attack and increase the yield.

## Acknowledgements

We thank Rosa María González-Martínez and Óscar Marín-Vázquez at University of Valencia (Spain) for their excellent help with insect rearing and laboratory management. We thank Marie-Christine François, Christelle Monsempès and Gabriela Caballero-Vidal at iEES-Paris (Versailles, France) for help in chemosensory gene annotation.

## Funding

This work has been supported by projects from the Spanish Ministry of Science, Innovation and Universities (No. AGL2014-57752-C2-2R and RTI2018-094350-B-C32). ALG was recipient of a PhD grant from the Spanish Ministry of Science, Innovation and Universities (No. BES-2015-071369). CMC was supported by a Juan de la Cierva contract (No. IJCI-2017-32501).

## Notes

https://www.ncbi.nlm.nih.gov/sra/PRJNA634227

### Competing Interest Statement

The authors have declared no competing interest.

