## Supplementary Figures for "Coupling transcriptomics and behaviour to unveil the olfactory system of *Spodoptera exigua* larvae"

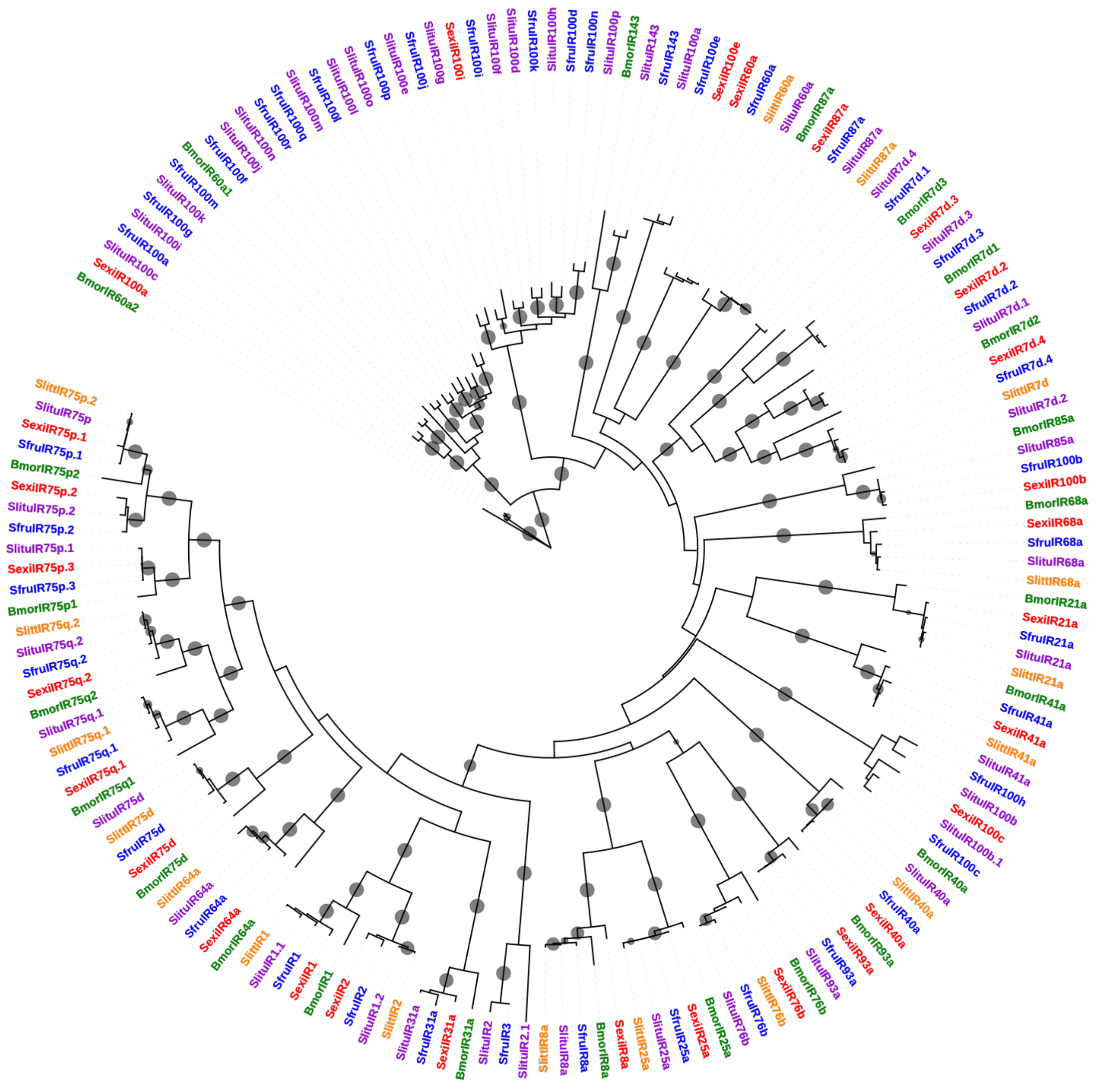


**Supplementary Figure 1. Phylogenetic tree of *Spodoptera exigua* ionotropic receptors (SexiIRs).** Maximum-likelihood (ML) tree built with protein sequences annotated from *S. frugiperda* (Gouin et al., 2017), *S. litura* (Zhu et al., 2018) and *B. mori* genomes (van Schooten et al., 2016) as well as putative proteins annotated from *S. littoralis* transcriptome (Walker et al., 2019).SexiIRs are shown in red, *S. frugiperda* IRs in blue, *S. litura* IRs in purple, *S. littoralis* IRs in yellow, and *B. mori* IRs in green. Grey dots show a bootstrap value higher than 80.

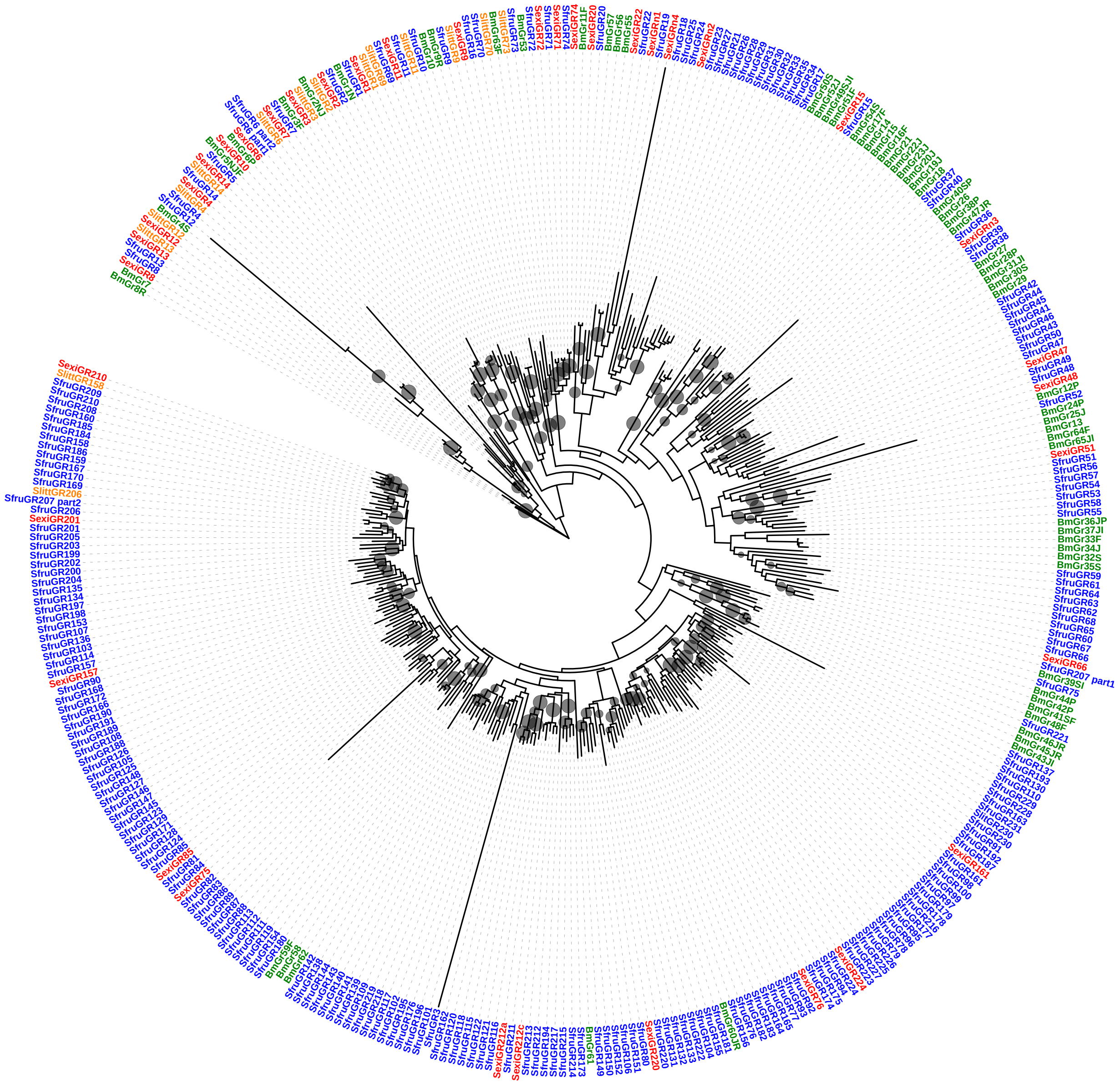

**Supplementary Figure 2. Phylogenetic tree of *Spodoptera exigua* gustatory receptors (SexiGRs).** Maximum-likelihood (ML) tree built with protein sequences annotated from *S. frugiperda* (Gouin et al., 2017) and *B. mori* genomes (Wanner and Robertson, 2008) as well as putative proteins annotated from *S. littoralis* transcriptome (Walker et al., 2019).SexiGRs are shown in red, *S. frugiperda* GRs in blue, *S. littoralis* GRs in yellow, and *B. mori* GRs in green. *S. exigua* amino acid sequences used for the tree are given in Supplementary Materials. Grey dots show a bootstrap value higher than 80.

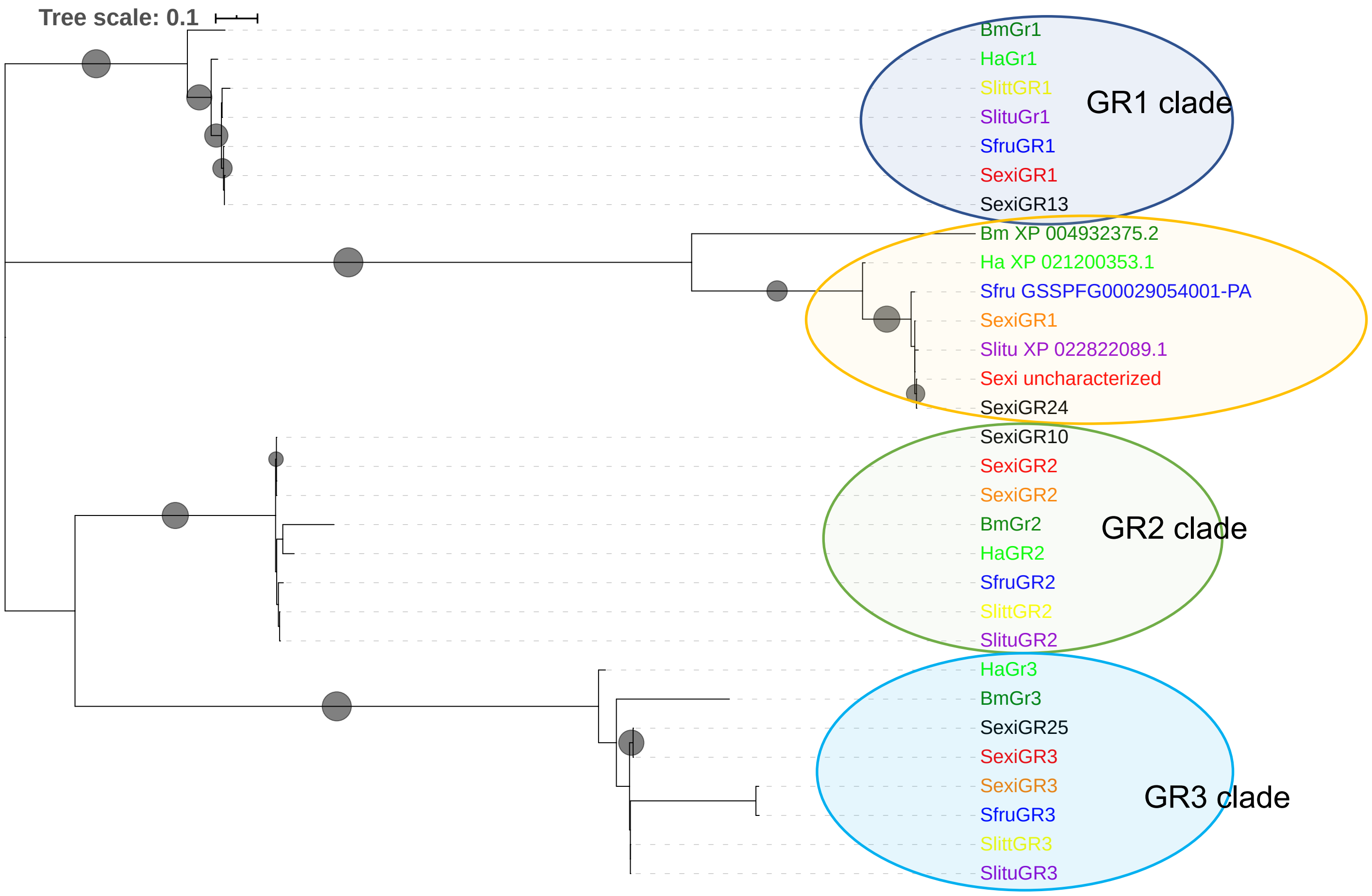


**Supplementary Figure 3. Phylogenetic tree of Lepidoptera $CO_2$ receptors.** Maximum-likelihood (ML) tree built with protein sequences of Lepidoptera $CO_2$ receptors. *S. exigua* sequences described in this study are shown in red, *S. exigua* GRs annotated by Zhang et al (2018) are shown in black, *S. exigua* GRs annotated by Du et al (2018) are shown in orange, *S. littoralis* sequences are in yellow (Walker et al., 2019), *S. frugiperda* sequences in blue (Gouin et al., 2017); *S. litura* genes in purple (Cheng et al., 2017), *B. mori* sequences in dark green (Wanner and Robertson, 2008), *H. armigera* genes in light green (Ning et al., 2016).

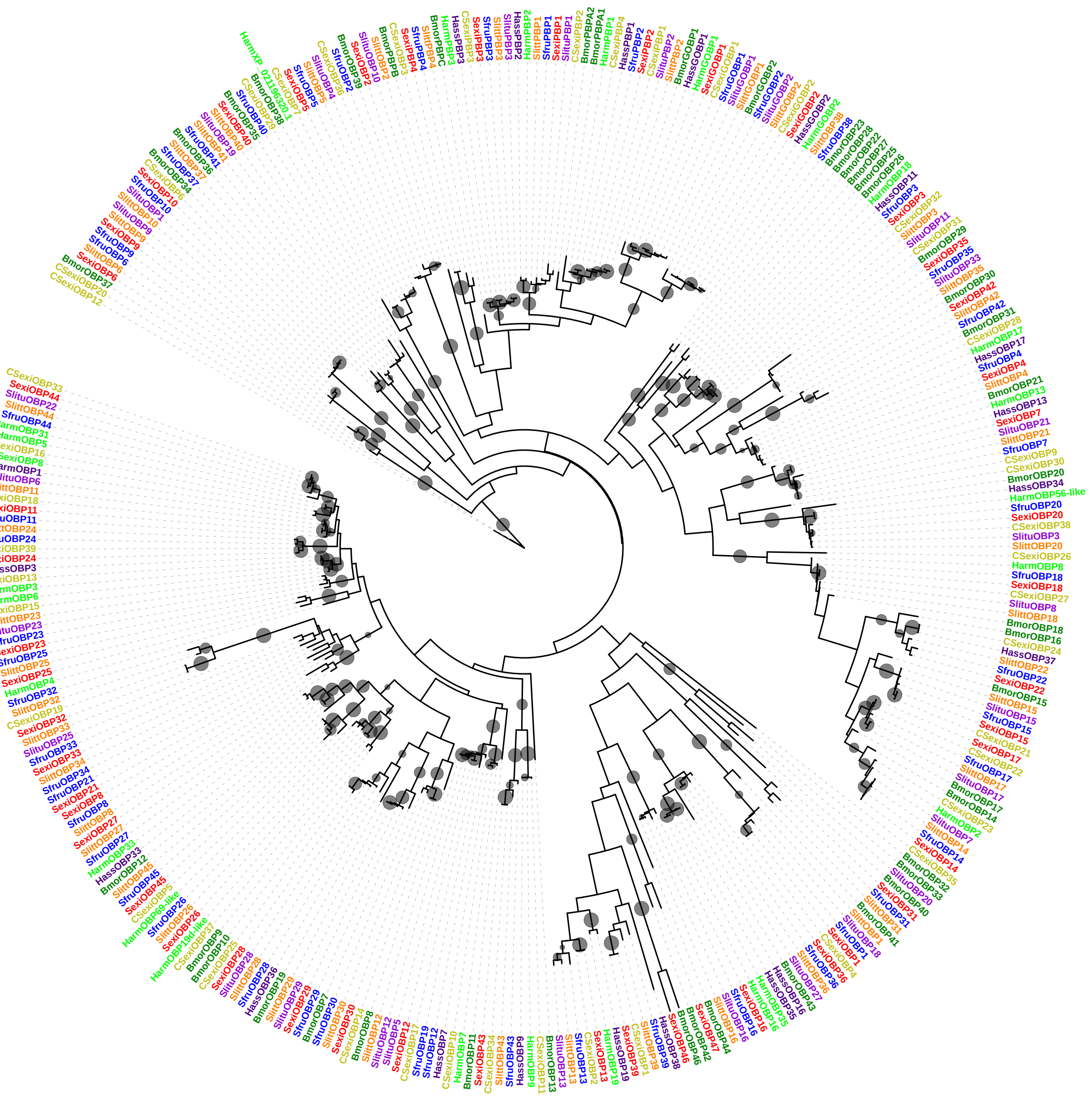

**Supplementary Figure 4. Phylogenetic tree of *Spodoptera exigua* odorant binding proteins (SexiOBPs).** Maximum-likelihood (ML) tree built with protein sequences annotated from *S. frugiperda* (Gouin et al., 2017) and *B. mori* genomes (Wanner and Robertson, 2008) as well as putative proteins annotated from *S. littoralis* (Walker et al., 2019), *S. litura* (Gu et al., 2015), *H. armigera* and *H. assulta* transcriptomes (Chang et al. 2017). SexiOBPs annotated in this study are shown in red whereas *S. exigua* OBPs annotated by Du et al (2018) in ochre, *S. frugiperda* OBPs in blue, *S. littoralis* OBPs in yellow, *B. mori* OBPs in dark green, *S. litura* OBPs in light purple, *H. armigera* OBPs in light green, *H. assulta* OBPs in dark purple. Grey dots show a bootstrap value higher than 80.

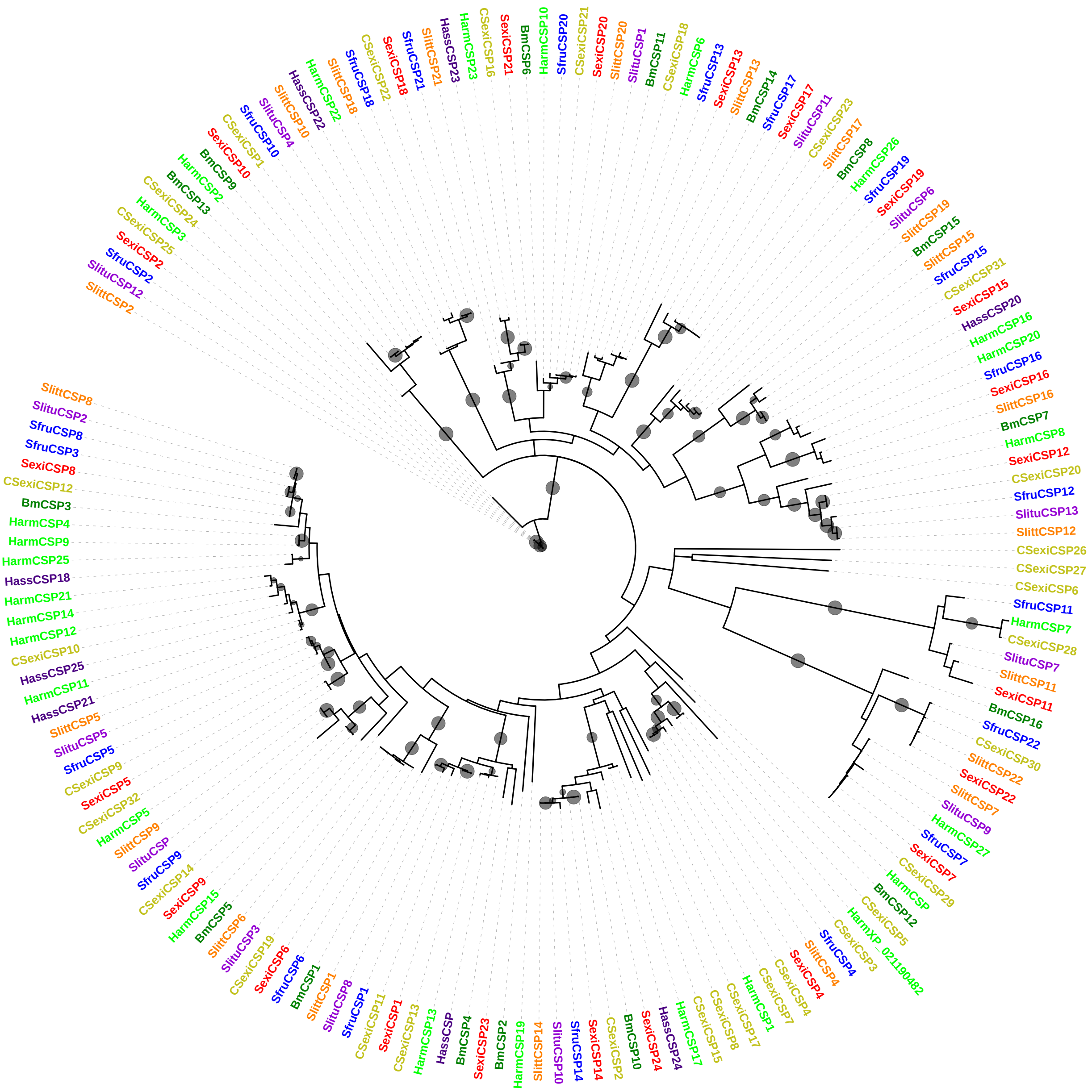


**Supplementary Figure 5. Phylogenetic tree of *Spodoptera exigua* chemosensory proteins (SexiCSPs).** Maximum-likelihood (ML) tree built with protein sequences annotated from *S. frugiperda* (Gouin et al., 2017) and *B. mori* genomes (Wanner and Robertson, 2008) as well as putative proteins annotated from *S. littoralis* (Walker et al., 2019), *S. litura* (Gu et al., 2015), *H. armigera* and *H. assulta* transcriptomes (Chang et al. 2017). SexiCSPs annotated in this study are shown in red whereas *S. exigua* CSPs annotated by Du et al (2018) (CSexiCSPs) in ochre, *S. frugiperda* CSPs in blue, *S. littoralis* CSPs in yellow, *B. mori* CSPs in dark green, *S. litura* CSPs in light purple, *H. armigera* CSPs in light green, the *H. assulta* CSPs in dark purple. Grey dots show a bootstrap value higher than 80.

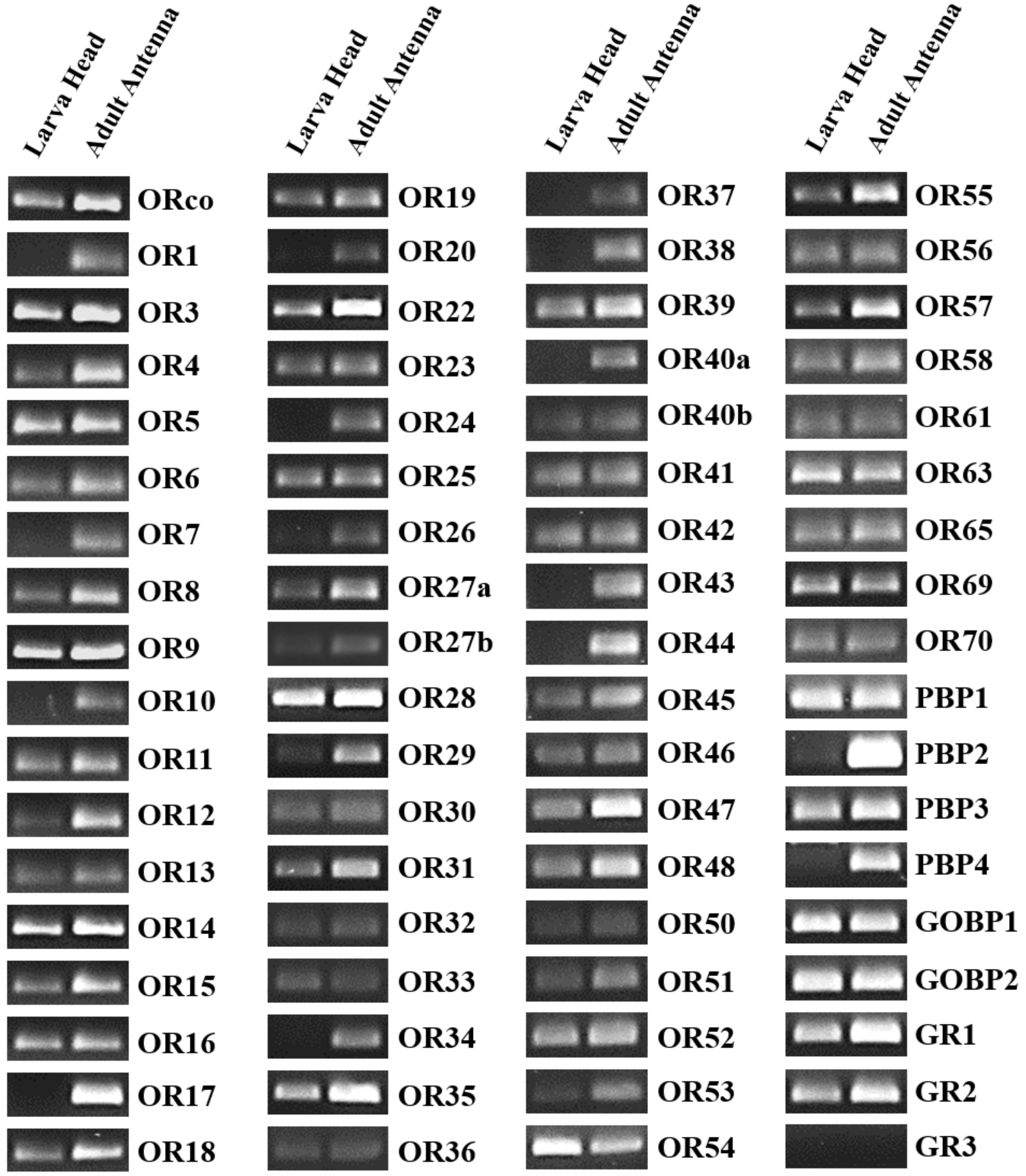


**Supplementary Figure 6. Expression profiles of odorant receptors and selected odorant biding proteins and gustatory receptors in larvae and adults.**

RT-PCR assays were performed using gene specific primer pairs and cDNAs from different *Spodoptera exigua* tissues: larva head and adult antenna. PCR products were analysed on 2% agarose gels. ORs: odorant receptors, PBP: pheromone binding proteins, GOBP: general odorant binding proteins, GRs: gustatory receptors.

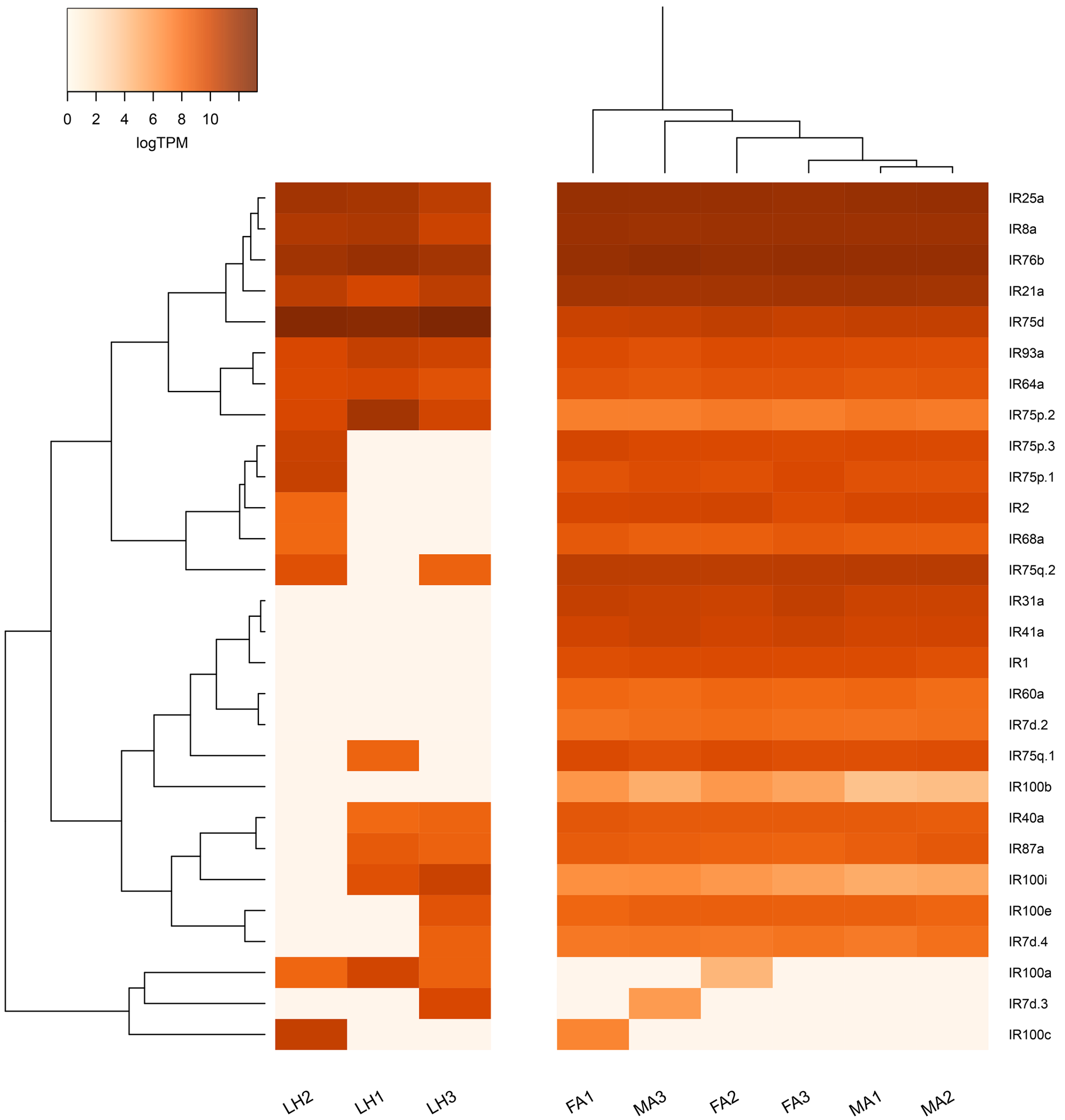


**Supplementary Figure 7. Heat-plot of ionotropic receptors (SexiIRs) expression in the head of *Spodoptera exigua* larvae and adult antennae.** Colour plots represent Log2 of transcripts per million (TPM) values estimated by RSEM. Light orange colours indicate low expression and dark orange ones indicate high expression. LH: Larvae Head. MA: Male Antennae. FA: Female Antennae.

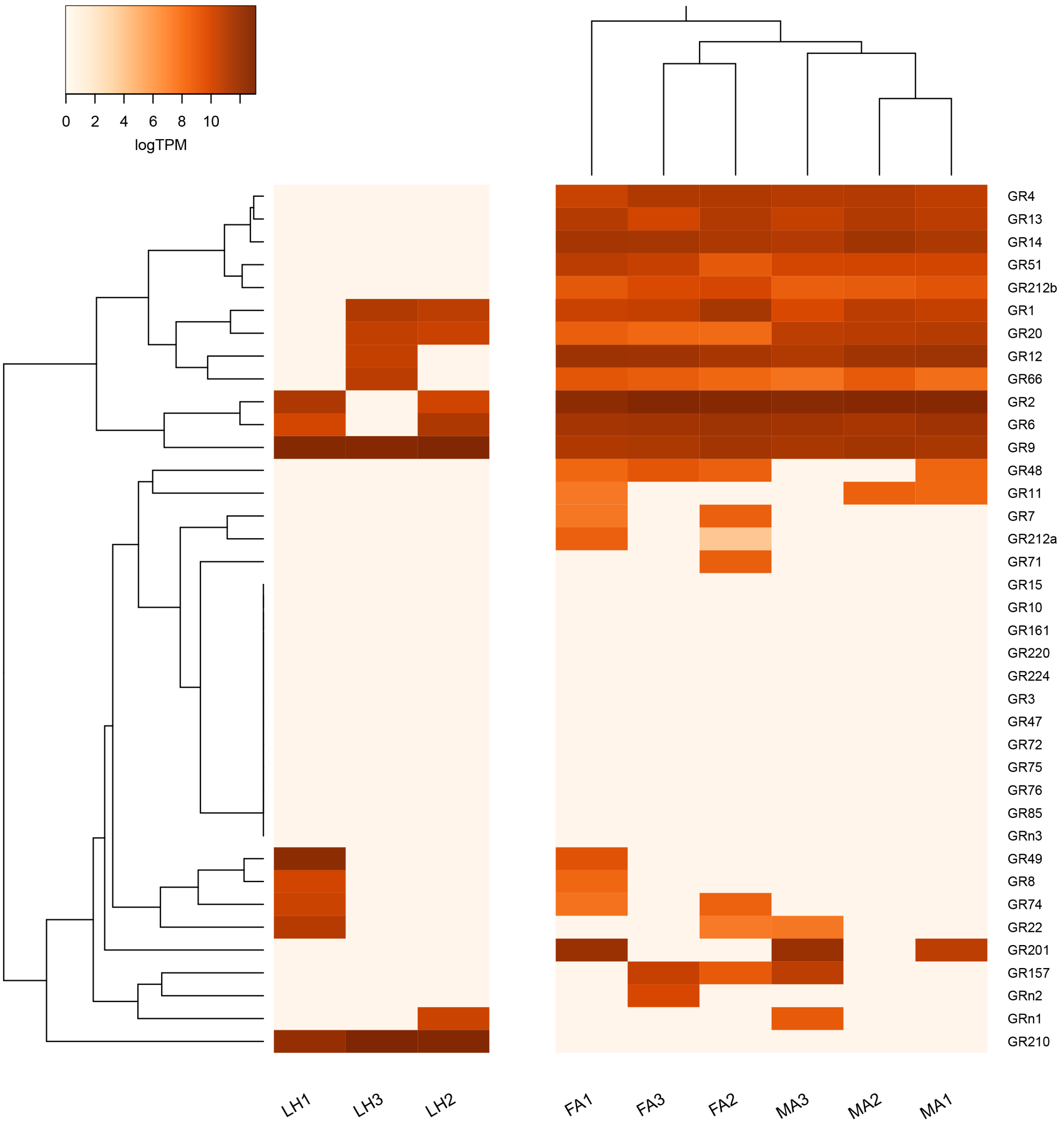


**Supplementary Figure 8. Heat-plot of gustatory receptors (SexiGRs) expression in the head of *Spodoptera exigua* larvae and adult antennae.** Colour plots represent log2 of transcripts per million (TPM) values estimated by RSEM. Light orange colours indicate low expression and dark orange ones indicate high expression. LH: Larvae Head. MA: Male Antennae. FA: Female Antennae. Asterisks indicate statistically significant differences between male and female antenna samples identified by EdgeR analysis (FDR<0.05).

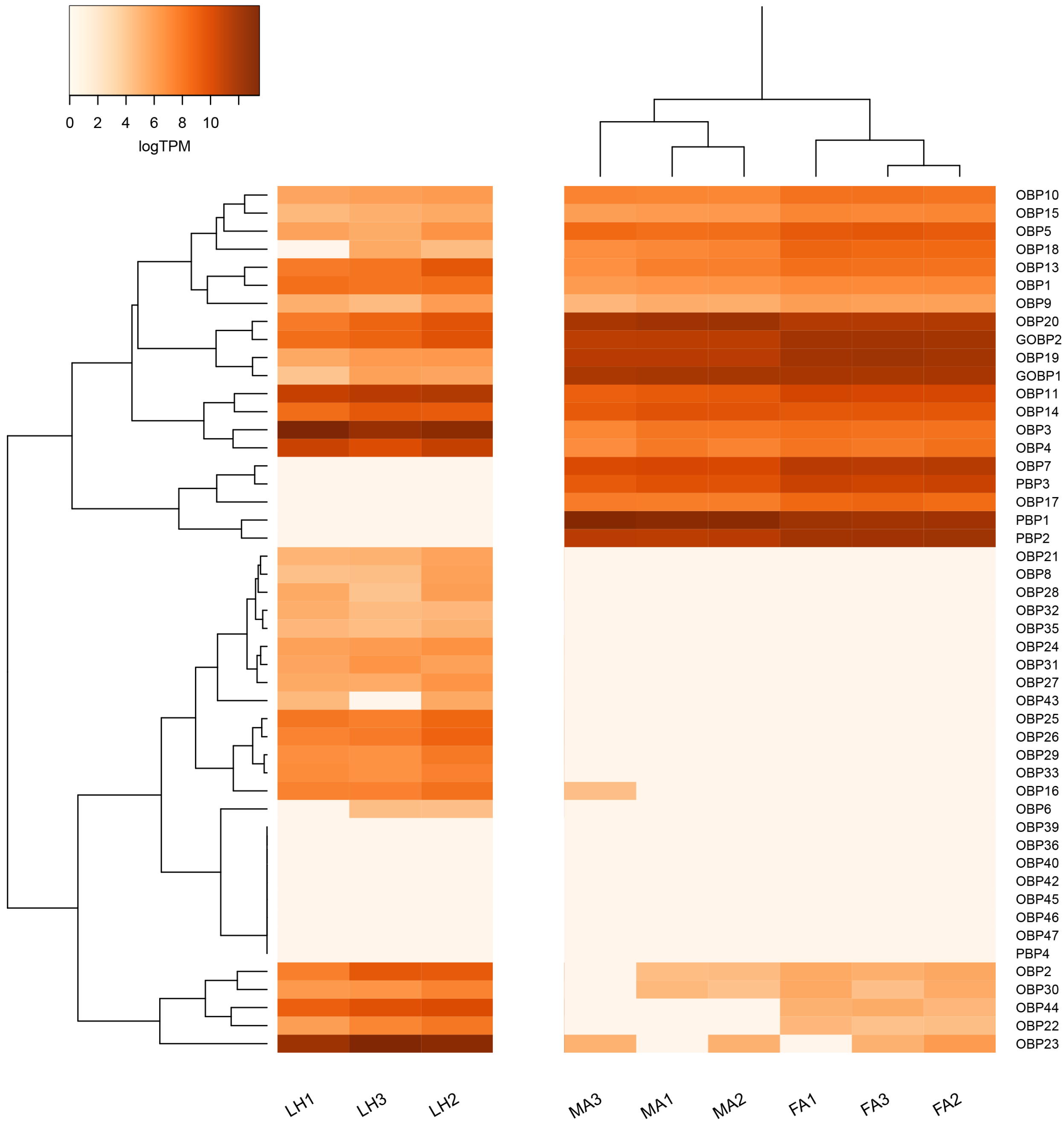


**Supplementary Figure 9. Heat-plot of odorant binding proteins (SexiOBPs) expression in the head of *Spodoptera exigua* larvae and adult antennae.** Colour plots represent log2 of transcripts per million (TPM) values estimated by RSEM. Light orange colours indicate low expression and dark orange ones indicate high expression. LH: Larvae Head. MA: Male Antennae. FA: Female Antennae. Asterisks indicate statistically significant differences between male and female antenna samples identified by EdgeR analysis (FDR<0.05).

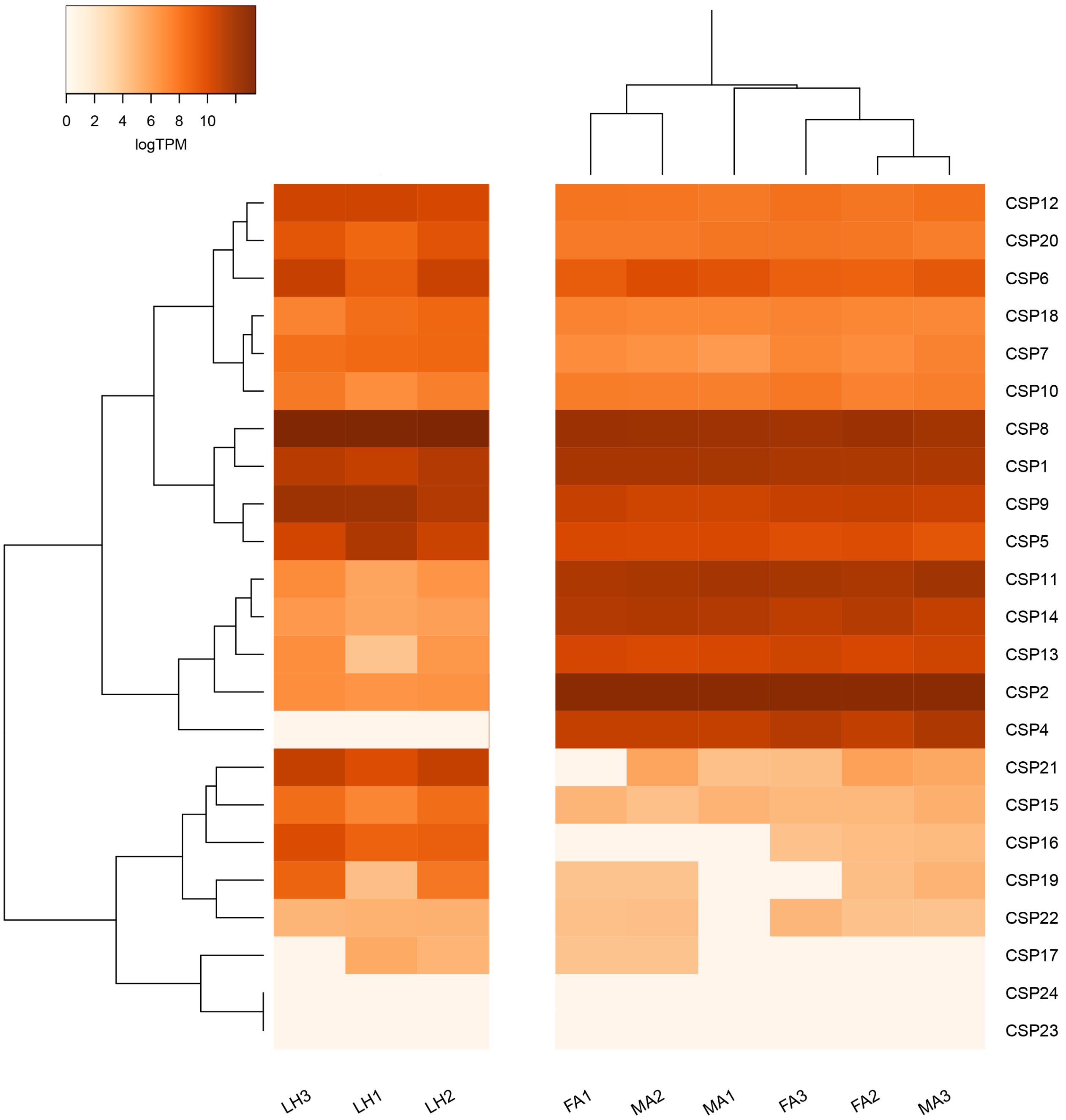


**Supplementary Figure 10. Heat-plot of chemosensory proteins (SexiCSPs) expression in the head of *Spodoptera exigua* and adult antennae.** Colour plots represent log2 of transcripts per million (TPM) values estimated by RSEM. Light orange colours indicate low expression and dark orange ones indicate high expression. LH: Larvae Head. MA: Male Antennae. FA: Female Antennae. Asterisks indicate statistically significant differences between male and female antenna samples identified by EdgeR analysis (FDR<0.05).
